# Assessing *Arthrospira platensis* Growth Rates in three Jeju Local Medium, Including Jeju Magma Seawater

**DOI:** 10.1101/2023.10.30.564724

**Authors:** Huey Jang

## Abstract

Jeju Magma Seawater (JMS), naturally formed beneath the bedrocks of Jeju island, consists of high micronutrient concentrations such as Zinc (Z, 0.019 mg/L), Iron (F, 0.015 mg/L), Manganese (M, 0.008 mg/L), Vanadium (V, 0.015 mg/L), Selenium (S, 0.013 mg/L) and Germanium (G, 0.002 mg/L). Despite the extensive research conducted upon the conventional Zarrouk medium used in Arthrospira platensis culture industry, the commercial potential of JMS on the growth of Arthrospira platensis has been poorly investigated. With respect to growth rate and cost efficiency, the purpose of the study is to discern and compare the growth of Arthrospira platensis in JMS with other accessible mediums on Jeju island. This study investigated the effects of three different growth mediums: Jeju Magma Seawater (JMS), Jeju seawater (JS), and Artificial seawater (AS) to assess their impact on the growth of Arthrospira platensis measured by biomass concentration (g L-1). The ANOVA results showed that JMS exhibited higher growth levels compared to JS ((p = 0.001) < 0.05) and AS (p=0.001). The results of the t-test revealed a statistically significant disparity between mean mass growth in JMS micronutrient-present environment and JMS micronutrient-absent environment (T = 7.94, p (= 4.57e-05) < 0.05). However, based on the growth rate over this experiment, it is estimated that the cost of cultivation required to obtain the same amount of mass is more than seven times higher in JMS compared to conventional Zarrouk medium. Therefore, it is believed that there are limitations to immediate commercialization.

## 1. Introduction

*Arthrospira platensis*, is cultivated on a scale of 12,000 metric tons per annum for both commercial and environmental uses (Masojídek *et al*., 2014): for example additional food supplements for humans (Aly *et al*., 2023), animals (European commision., 2022), and aquaculture or ‘liquid trees’ where 1kg (dry mass) of *Arthrospira platensis* consumes 1.83kg of CO_2_ against climate change (Ravelonandro *et al*., 2010). *A.platensis* is a planktonic filamentous cyanobacterium composed of individual cells (Masojídek *et al*., 2008), and is the most cultivated photosynthetic prokaryote due to its versatility. The biomass contains rich amounts of proteins (∼60%) and carbohydrates(∼15%), lipids, phycobiliproteins, etc (Torzillo *et al*., 2008). Apart from its advantageous functions, the species also has garnered interest due to its ability to thrive in poor-quality water, like wastewater in alkaline environments, provided there is an ample supply of nutrients and sunlight (Hena *et al*., 2017). Hence, it is relatively easy to cultivate, as it can adapt to different water environments with various salinities: freshwater, brackish lakes, and alkaline saline lakes (Vonshak A *et al*., 1997). Due to its versatility and availability, the mass cultivation of *A.spirulina* is expected to expand (Tzachor *et al*., 2022).

In terms of artificial culture setting, the two crucial factors toward the growth of *Arthrospira platensis* are: environmental factors (lighting, temperature, inoculation volume, turbulence, etc.) and the Medium (Alfadhly *et al*., 2022). The optimum values of environmental factors have been extensively experimented and established upon (Kebede *et al*., 1996). Likewise, an established type of culture medium, the Zarrouk medium (Zarrouk *et al*., 1966), has been used since 1966 for the cultivation of *A.platensis*.

However there are definite economical drawbacks to the usage of the Zarrouk medium. The overall cost of the formation of Zarrouk medium covers approximately 35% of the total cost of biomass production (Costa *et al*., 2019). The financial loss is not only due to its manufacturing costs, but also due to its inability to be recycled. The Zarrouk medium loses its productivity when it is reused for another batch of culture due to secondary metabolites, non-consumed nutrients and cellular debris that accumulate. This changes the properties of the solution, hence impairing the optimum conditions set for maximum growth. Increased alkalinity creates uncertainty of nutrient concentration, while microorganisms and dissolved organic matter (DOM) can negatively affect the next batch (Loftus *et al*., 2017). Research shows that compared with the unused medium, the growth rate of microalgae substantially decreases with each time the medium is recycled (Yuan *et al*., 2019). If a medium with lower cost per unit compared to the growth rate of algae can be established, it can reduce the economic expenses for the medium, thus increasing its industrial utility value.

Considering these difficulties posed by the use of the Zarrouk medium, researches continuously reveal novel types of mediums with different compositions of micronutrients and financial demands for both formation and reusal (Raoof *et al*., 2006).

Jeju Magma Seawater (JMS) is produced when seawater runs underground and is naturally filtered through volcanic bedrock. Because volcanic bedrock is formed of basalt, its pores allow sea water to drain inwards underground and is most abundantly found in the eastern part of the island as it has most of the volcanic bedrock which seawater can easily run through. Because it originates from sea water it is not only renewable but also naturally saline providing a compatible salinity condition for *Arthrospira platensis* which thrives in salinities ranging from 30-40 ppt (Piu *et al*., 2022). JMS also consists of high micronutrient concentrations, as much as 10 times compared with natural seawater. One of the unique properties of JMS is the high concentrations of inorganic components: Zinc (Z, 0.019 mg/L), Iron (F, 0.015 mg/L), Manganese (M, 0.008 mg/L), Vanadium (V, 0.015 mg/L), Selenium (S, 0.013 mg/L) and Germanium (G, 0.002 mg/L), of which its effect on the growth of *Arthrospira platensis* is yet to be researched upon. With the readily set composition of micronutrients and salinity, minimum prior treatment will be required to use JMS as a medium for algae, which significantly reduces the manufacturing cost compared to the widely spread Zarrouk medium.

Compared to the Zarrouk medium, JMS is not artificially composed of, but is naturally produced. Hence, the widespread utilization of JMS has the potential to accelerate the mass production of *Arthrospira platensis* without generating substantial cost. There are demands to investigate the effects of JMS as an more accessible alternative to Zarrouk medium (Kim *et al*., 2019, Lee *at al*. 2018ab, 2019). With a similar salinity and pH to the Zarrouk medium, JMS has the potential to provide a suitable environment for *Arthrospira platensis* cultures (Lee *et al*., 2018a). In a previous research, micronutrients extracted from JMS showed a potential to support the growth of species *Spirulina maxima*, another industrious algae species closely related with *A. platensis* (Lee *et al*., 2017, Lee *et al*., 2018ab).

In this research, JMS’s effects on growth of *Arthrospira platensis* was compared against two other local mediums in Jeju Island: Jeju Seawater and Artificial Seawater, thus forming the research question - To what extent do three different mediums: Jeju Magma seawater (JMS), Jeju seawater (JS), and Artificial seawater (AS) affect the growth of the *Arthrospira platensis’* biomass concentration (g L^-1^).

## 2 Methods and Materials

### (1) Medium Collection

JMS was provided by Jeju Korea Institution of Ocean Science and Technology (KIOST Jeju). Artificial seawater was formed by dissolving 39g of reef salt (containing monir chlorine, sodium, magnesium, sulfur, calcium, potassium, boron and strontium) with 1L of water, which created a salt solution of 33psu. Jeju seawater from the Jeju coastline was collected from the coordinates: 33.23252° N, 126.31371° E on August.27. It was expected to contain carbon (TC), dissolved inorganic nitrogen (DIN), dissolved inorganic phosphorus (DIP) and dissolved silicon (DSi) in addition to the components of AS (Moon *et al*., 2020)).

### (2) Methods opted for Measurement

All strains of *A.platensis* were distributed from the Korean Collection for Type Cultures (KCTC). Spectrophotometry with O.D_750_ was used to determine the density and mass of the specimen throughout 2 weeks of cultivation. The measured value using O.D_750_ is known to have a positive correlation with biomass concentration (g/L), Chlorophyll content, Co2 fixation, and cell number. Thereby, results obtained through this research are expected to be able to be converted and compared with data from prior studies and literature. Growth rate was obtained by the difference between the initial mass and the mass cultured over a period of two weeks, divided by the initial mass.”

### (3) Experiment Procedure

5 sets of 3 containers were filled up with 300ml of each type of medium: JMS, AS, and JS with a total of 15 containers. A rubber tube with a 5mm diameter connected to an air pump was placed inside each container. Then, *Arthrospira platensis* culture (1×10^6 cells/L) was suspended in a total of 250 ml of distilled water and was equally divided into 15 culture samples of 16.5ml using a pipette. Each sample was added to the container.

All batches received light from light stands of the same model. Regarding prior studies, they were exposed to continuous light for 24hrs, due to the self shading effect (Ravelonandro *et al*., 2011, W. K. Lee pers. Comm. May 13th). Meanwhile, in order for the culture to be equally distributed across the containers agglomeration had to be prevented (Gibson *et al*., 1990). The method opted was a bubble column, as it was proven to be the most effective method of turbulence proven by Richmond et al. (Richmond *et al*., 1996).

From each batch, 6 measurements were taken per day. Before each measurement, any agglomeration of algae was scraped from the surfaces using a spatula, and dispersed using the pipette. When the solution had been mixed thoroughly, 1ml sample was taken directly from the container using a pipette, and was transferred to a cuvette. Before the algae sedimented to the bottom, it was quickly placed in a spectrophotometer, and the optical density at 750 nm (O.D_750_) was measured. This process was repeated 6 times per container, and was recorded on a data sheet.

### (4) Statistical Analysis

ANOVA test was used as the data was consecutive (not ranked) and there were more than two groups. The Tukey test was employed to assess whether there were significant differences between each pair of groups. t-test was used to assess the significance of the impact of minerals that only consist in JMS on the growth of *A. platensis*. Hence, it was conducted between two groups: JMS, and the other group without the certain minerals (JS and AS).

## 3 Results and Discussion

### (1) Measurement of Algae Growth in Different Environments

To compare the growth of algae in three different environments, namely JMS, Jeju seawater (JS), and Artificial seawater (AS), Optical density at 750 nm (O.D_750_) was measured and converted this value to biomass concentration (gL^-1^) and chlorophyll content (gL^-1^). The results showed that the average values of O.D_750_, biomass concentration (gL^-1^), and chlorophyll content (gL^-1^) were highest in the JMS condition, followed by AS and then JS. In terms of O.D_750_, JMS had a value of 0.42, which was higher than AS by 0.23 and higher than JS by 0.22. Additionally, chlorophyll content (gL^-1^) followed the similar trend to O.D_750_, but after a couple of cultivation days, the number of cells in the AS condition exceeded that in the JS condition. JS consistently had the fewest cells across all time points. biomass concentration (gL^-1^) also exhibited a similar trend to O.D_750_ and the chlorophyll content (gL^-1^), reaching the threshold in JMS algae first, showing similar trends in growth.

### (2) Statistical Comparison of Algae Growth

First, an analysis of variance (ANOVA) was conducted to assess the differences in the mean growth rate among three distinct mediums: JSM, JS, and AS. This ANOVA aimed to determine whether there were statistically significant variations in growth level across these environments. Figure 1 presents the statistics for the growth rate in each medium (F = 7.92, p (= 0.006) < 0.05). The calculated F-value exceeded the critical value, suggesting that at least one of the group means significantly differs from the others. Furthermore, Post-hoc comparisons were performed using the Tukey-Kramer method to determine which specific pairs of environments exhibited statistically significant differences in the level. Results revealed that JMS exhibited significantly higher growth levels compared to JS (p = 0.01) and AS (p = 0.01). No significant differences were observed between JS and AS (p = 0.98). Mean difference between JS and AS was -0.01, and the standard deviations were -0.1587 and 0.1386.

**Fig. 1.**
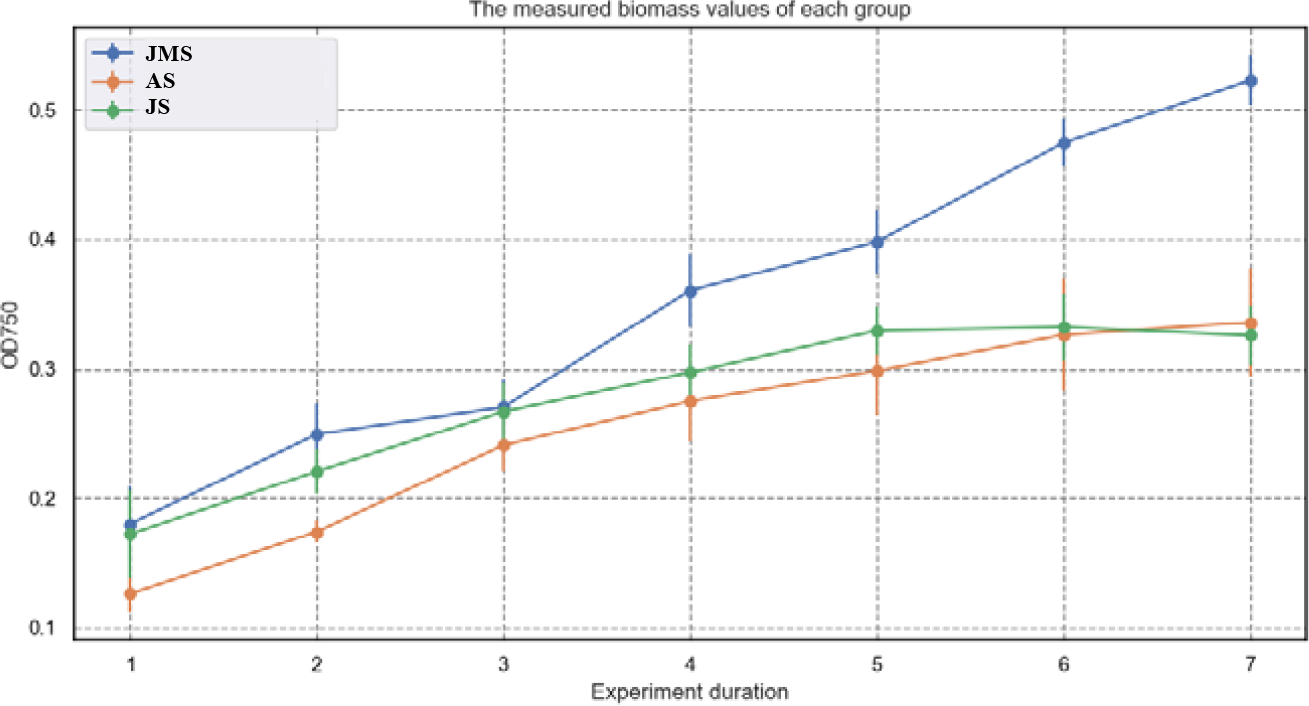
Mean biomass concentration (gL^-1^) during the three times of *Arthrospira platensis* cultivations from different three local medium: JMS (Jeju magma seawater), AS (Artificial seawater) and JS (Jeju seawater). Bars represent 95% confidence intervals.

Secondly, It has been established that Zinc (Z, 0.019 mg/L), Iron (F, 0.015 mg/L), Manganese (M, 0.008 mg/L), Vanadium (V, 0.015 mg/L), Selenium (S, 0.013 mg/L) and Germanium (G, 0.002 mg/L), potent catalysts for algae growth, is exclusively present in JMS. To assess whether there is a significant average difference in growth rates between the unique substance-containing JMS and other environments lacking it (JS, AS), it was tested whether the observed discrepancies in the means of the sample groups are attributable to chance or if they indeed signify a genuine distinction between the populations (Figure 2). The mean O.D_750_ for the medium with components was 4.0, accompanied by a standard deviation of 0.05. Similarly, for the medium without component, the mean score was 1.7, along with a standard deviation of 0.03. The outcomes of the t-test reveal a statistically significant disparity between the mean scores of the substance-present environment and the substance-absent environment (t = 7.94, p (= 4.57e-05) < 0.05).

**Fig. 2.**
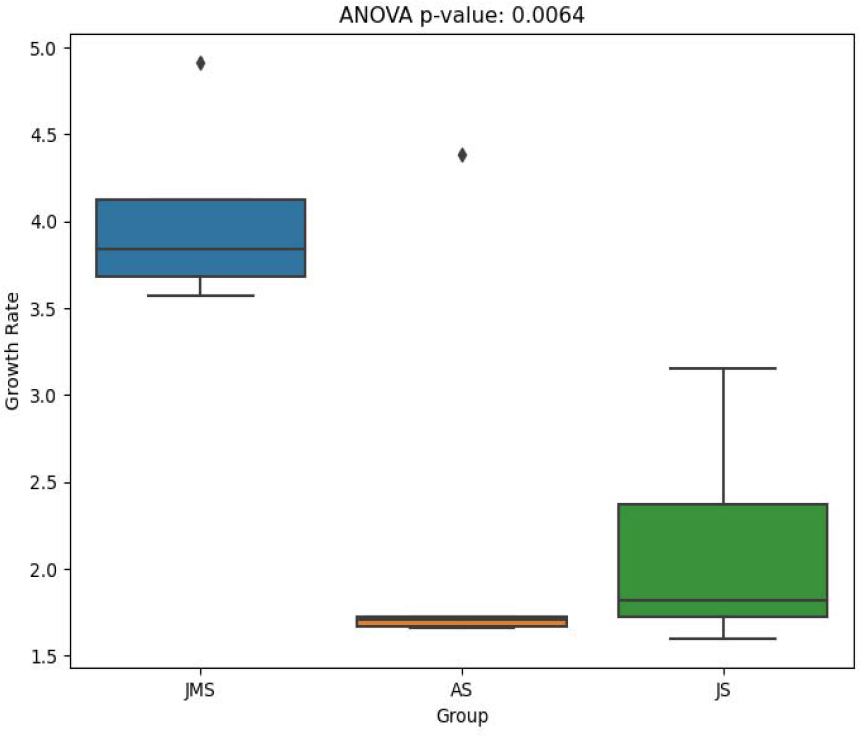
Comparison on the *Arthrospira platensis*’ growth rate between the three different mediums (Jeju magma seawater (JMS), Artificial seawater (AS), Jeju seawater (JS)). Bars represent 95% confidence intervals.

**Fig. 3.**
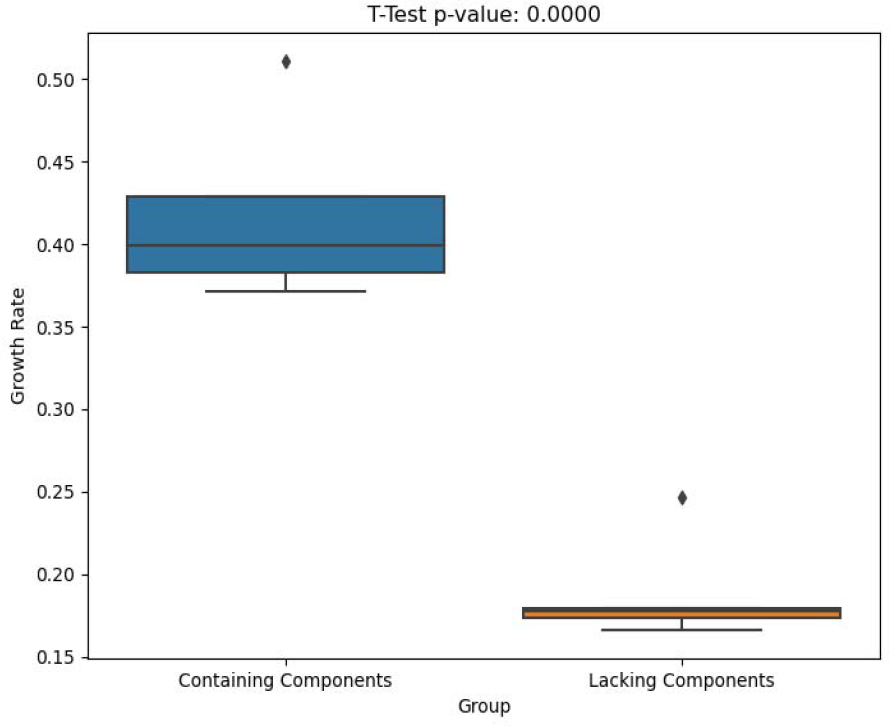
Comparison on *Arthrospira platensis*’ growth rate between with and without Zinc (Z, 0.019 mg/L), Iron (F, 0.015 mg/L), Manganese (M, 0.008 mg/L), Vanadium (V, 0.015 mg/L), Selenium (S, 0.013 mg/L) and Germanium (G, 0.002 mg/L) components. Bars represent 95% confidence intervals.

**Fig. 4.**
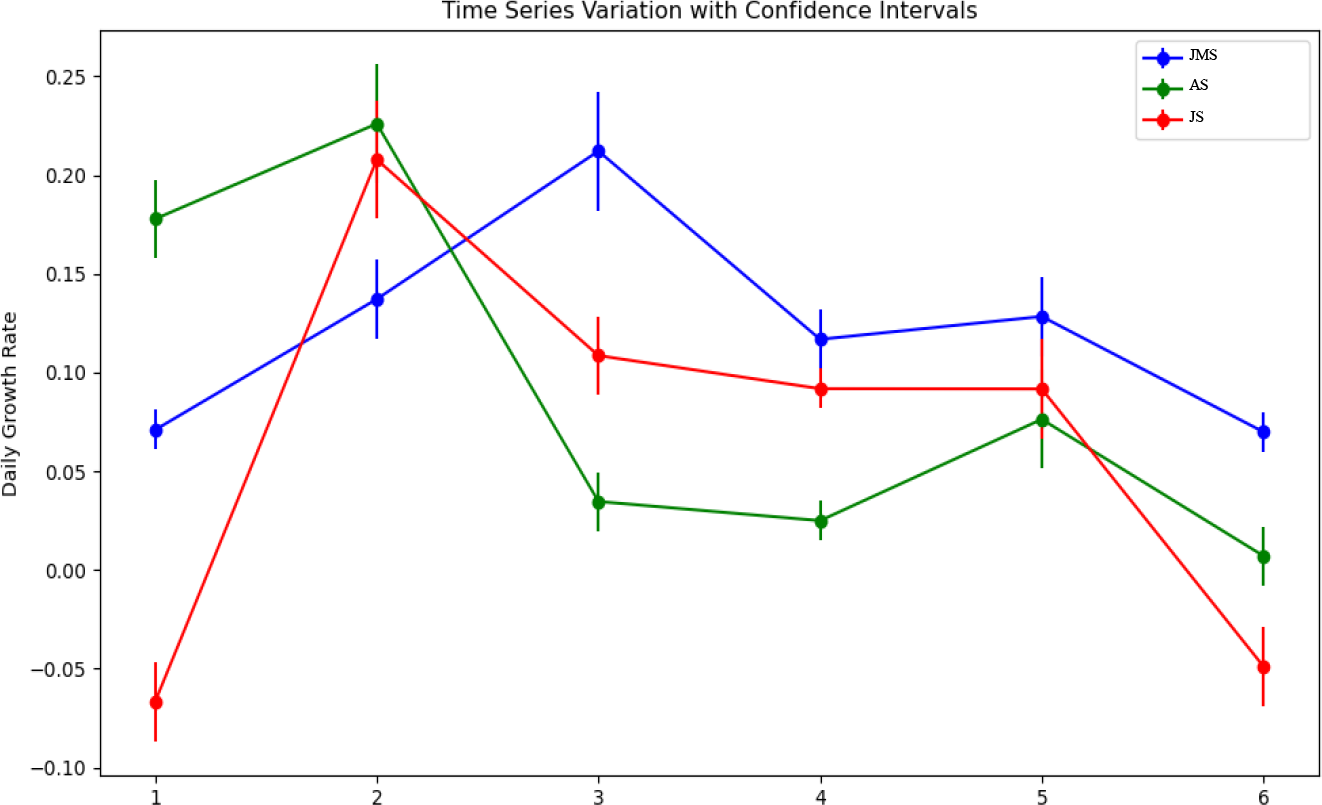
Daily changes in *Arthrospira platensis*’ growth rates between the three different mediums (Jeju magma seawater (JMS), Artificial seawater (AS), Jeju seawater (JS)). Bars represent 95% confidence intervals.

### (3) Temporal Variation in Algae Growth Rates Across Different Environments

This study examined the changing growth rates of the algae over five repetitions in three different environments to study how each medium influences the growth patterns of the target algea. These data would be valuable for future modeling studies and determine the medium replacement cycle required for the future commercialization of JMS by providing the slope of daily growth rate and peak growth rate.

In all experiments, the growth rate of JMS consistently reached the highest peak and continued to increase the latest. On the other hand, the deceleration of the growth rate was slower compared to the other two media, and it did not drop below zero until the end of the experiment. In JMS, the algae growth rate exhibited an linear trajectory during the first 3 days. However, the growth rate fluctuated and decreased towards day 7. In AS, the growth rate showed a similar trend to that of JMS, where the growth rate increased during the initial 2 days, and steadily declined reaching plateau on day 5, and decreasing sharply afterwards reaching near 0. JS also showed a similar pattern of an increase followed by a steady decrease of growth rate, but it is unique as all sample cultures showed a negative growth rate during the last two consecutive days.

### (4) Economic Comparison with Zarrouk medium Based on Results

Based on the measured growth rates, the cost-effectiveness of JMS was evaluated by comparing the unit input costs with the pre-established medium in industry sector, Zarrouk medium. Through a comparison of costs, the economic viability of JMS was assessed. From previous research, the inferred market price per unit kg of algae produced was $8.43. Previous research that utilised the Zarrouk medium to cultivate *Arthrospira platensis* in a similar experimental condition with this study showed a growth rate of OD_750_ 16 per day (Lim *et al*., 2020). The cost of the Zarrouk medium per the unit growth was determined to be $0.12. Thus, the profit balance when cultivating algae using Zarrouk medium in a typical industrial setting follows the formula: (OD_750_ 1 + OD_750_ 16)($8.43-$0.12) = 141.27 ($*OD_750_). In contrast, in this study, the unit growth rate of JMS was determined to be OD_750_ 2.06, and its unit cost was set at $0—this is a cost that does not include transportation expenses, and since only the cost of materials required for the synthesis of Zarrouk medium was considered, it allows for a rough comparison. Consequently, the derived revenue formula when utilising JMS for algae cultivation is: $8.43(OD_750_ 1+ OD_750_ 2.06)k = 25.80 ($*OD_750_). Hence, it could be the potential conclusion that JMS might yield only a limited profit, potentially making commercialization challenging, without further research that enables an economic evaluation of the fact that JMS has almost no environmental impact in the purification and discharge processes.

## 4. Conclusion

In this research, we explored how various growth mediums affect the development of Arthrospira platensis, a widely farmed cyanobacterium with a range of practical uses. This study aimed to find alternatives to the Zarrouk medium, which has raised concerns regarding the environmental and economic sustainability in cultivating this microorganism. To achieve this, the study focused on testing and comparing the growth performance of three local mediums from Jeju Island: Jeju Magma Seawater (JMS), Jeju seawater (JS), and Artificial Seawater (AS), aiming to understand their influence on the maximum growth of Arthrospira platensis as measured by a spectrophotometer (O.D_750_). Jeju Magma Seawater (JMS) consistently showed higher growth mass for Arthrospira platensis compared to Jeju seawater (JS) and Artificial Seawater (AS). Also, JMS’s components, including Zinc, Iron, and others, contributed to its enhanced growth compared to the other two mediums. In terms of temporal growth, all three mediums initially displayed rapid growth for the initial days, followed by a slowdown after three days. The growth rate increase of JMS was sustained longer than other mediums, resulting in the peak occurring later and also exhibiting the slowest rate during decline phase. While JMS offers growth advantages, economic analysis revealed it might generate 6 times lower profit than the Zarrouk medium, posing challenges for commercial viability.

